# Design of an epitope-based peptide vaccine against *Cryptococcus neoformans*

**DOI:** 10.1101/434779

**Authors:** Isra Khalil, Ibtihal Omer, Islam Zainalabdin Abdalgadir Farh, Hanaa Abdalla Mohamed, Hajr Abdallha Elsharif, ALazza Abdalla Hassan Mohamed, Mawadda Abd-Elraheem Awad-Elkareem, Mhamed Ahmed Salih

## Abstract

**Introduction:** This study aimed to design an immunogenic epitope for Cryptococcus neoformans the etiological agent of cryptococcosis using in silico simulations, for epitope prediction, we selected the mannoprotein antigen MP88 which it’s known to induce protective immunity.

**Material & method:** A total of 39 sequences of MP88 protein with length 378 amino acids were retrieved from the National Center for Biotechnology Information database (NCBI) in the FASTA format were used to predict antigenic B-cell and T cell epitopes via different bioinformatics tools at Immune Epitope Database and Analysis Resource (IEDB). The tertiary structure prediction of MP88 was created in RaptorX, and visualized by UCSF Chimera software.

**Result:** A Conserved B-cell epitopes **AYSTPA, AYSTPAS, PASSNCK, and DSAYPP** have displayed the most promising B cell epitopes. While the **YMAADQFCL, VSYEEWMNY** and **FQQRYTGTF** they represent the best candidates T-cell conserved epitopes, the 9-mer epitope **YMAADQFCL** display the greater interact with 9 MHC-I alleles and HLA-A*02:01 alleles have the best interaction with an epitope. The **VSYEEWMNY** and **FQQRYTGTF** they are non-allergen while **YMAADQFCL** was an allergen. For MHC class II peptide binding prediction, the **YARLLSLNA, ISYGTAMAV** and **INQTSYARL** represent the most Three highly binding affinity core epitopes. The core epitope **INQTSYARL** was found to interact with 14 MHC-II. The allergenicity prediction reveals **ISYGTAMAV, INQTSYARL** were non-allergen and **YARLLSLNA** was an allergen. Regarding population coverage the **YMAADQFCL** exhibit, a higher percentage among the world (69.75%) and the average population coverage was **93.01%.** In MHC-II, **ISYGTAMAV** epitope reveal a higher percentage (74.39%) and the average population coverage was (81.94%). This successfully designed a peptide vaccine against Cryptococcus neoformans open up a new horizon in Cryptococcus neoformans research; the results require validation by in vitro and in vivo experiments.

## Introduction

*Cryptococcus neoformans*, the predominant etiological agent of cryptococcosis **[1, 2].** Acquired by inhaling spores or desiccated yeast from the environment **[3, 4],** can cause life-threatening infections of the central nervous system in immunocompromised and immunocompetent individuals. Cryptococcal meningoencephalitis is the most common disseminated fungal infection in AIDS patients **[5–10]**, and a significant opportunistic infection in solid organ transplant recipients[7, 11, 12, 9] whereas exhibit the third most common invasive fungal infection among this group [13, 14] with a reported incidence of 1 to 5% and mortality of 20 to 40% [15, 16]. Despite therapy, mortality rates in these groups are high [7]. Globally 1 million cases of cryptococcal meningitis occur each year, resulting in approximately 625,000 deaths **[17, 18].**

Patients who survive Cryptococcal meningitis may have long-term sequelae related to their infection such as focal neurologic deficits, blindness, deafness, cranial nerve palsies and memory deficits, and may require prolonged therapy or experience disease relapses [15, 8].

Currently, cryptococcosis is treated with antifungal drugs, however, the use of antifungals is problematic because they are generally highly toxic, and have only a limited ability to eradicate an infection. Furthermore, excessive use of fungal drugs facilitates the emergence of resistant strains [19–23] for that a strong need for a vaccine against C. neoformans it became inevitable and it’s generally considered to be the most effective procedure of preventing infectious diseases.

The capsule is the most important virulence factor of the *Cryptococcus neoformans*. This structure consists of highly hydrated polysaccharides, It is also composed of mannoproteins (MPs) including 115, 98, 88, and 84 kDa were identified and characterized as *C. neoformans* immunoreactive antigens involved in the pathogenesis, and are potential cryptococcosis vaccine candidates **[24, 25].**

Strong evidence indicates that mannoproteins (MPs) from *C. neoformans* play a key role in inducing the T-cell mediated immune response, which is critical for antifungal protection [**25**–29], Also mannoproteins (MP) have the ability to elicit delayed-type hypersensitivity and Th type 1-cytokines, both critical to the clearance of this pathogenic yeast **[30–32].**

In this study, we aimed to design an immunogenic epitope for Cryptococcus neoformans using in silico simulations, for epitope prediction, we selected the mannoprotein antigen MP88 which it’s known to induce protective immunity.

Epitope-based vaccines are providing a rational alternative by improving immunity through the selection of only a subset of all epitopes capable of evoking the desired immune response. The b-cell epitope of a target molecule can be conjugated with a T-cell epitope to make it immunogenic. With the immunoinformatics, it is now possible to evaluate the immunogenic properties of proteins via computational methods (in silico) with high efficiency and confidence [33–35].

## Material and method

The flow chart demonstrates the overall procedures of peptide vaccine for *Cryptococcus neoformans* is illustrated in Figure1.

**Figure 1:**
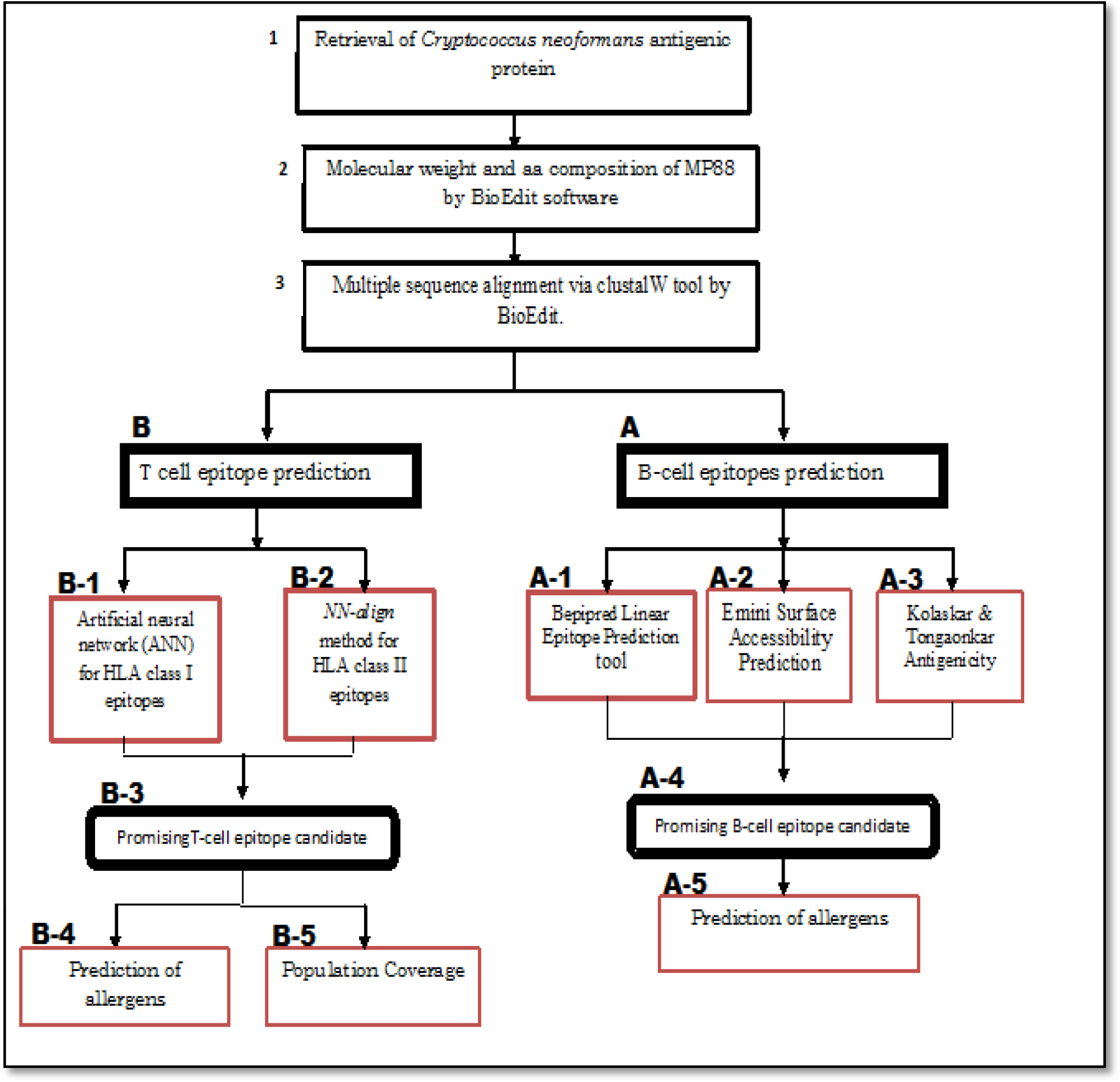
Graphical representation of Peptide vaccine design against *Cryptococcus neoformans*

### Retrieval of *Cryptococcus neoformans* antigenic protein

A total of 39 sequences of MP88 protein with length 378 amino acids were retrieved from the National Center for Biotechnology Information database (NCBI) (www.ncbi.nlm.nih.gov). All retrieved sequences belonged to genome sequencing and analysis program, Board Institute of MIT and Harvard, submitted in June 2017, In the FASTA format were further used for analysis. Molecular weight and amino acids composition of MP88 were determined by BioEdit software version 7.2.5. [Table 1], [Figure 2]. After that, multiple sequence alignment via clustalW tool was used to identify conserved regions in the protein sequences by using BioEdit.

**Table 1:** Protein: XP_012046953.1 immunoreactive mannoprotein MP88 [Cryptococcus neoformans var. grubii H99], Length = 378 amino acids, Molecular Weight = 38208.51 Daltons., illustrated by BioEdit software version 7.2.5

| Amino Acid |  | Number | Mol% |
| --- | --- | --- | --- |
| Ala | A | 48 | 12.70 |
| Cys | C | 8 | 2.12 |
| Asp | D | 14 | 3.70 |
| Glu | E | 15 | 3.97 |
| Phe | F | 11 | 2.91 |
| Gly | G | 44 | 11.64 |
| His | H | 4 | 1.06 |
| Ile | I | 12 | 3.17 |
| Lys | K | 3 | 0.79 |
| Leu | L | 20 | 5.29 |
| Met | M | 9 | 2.38 |
| Asn | N | 22 | 5.82 |
| Pro | P | 18 | 4.76 |
| Gln | Q | 12 | 3.17 |
| Arg | R | 6 | 1.59 |
| Ser | S | 53 | 14.02 |
| Thr | T | 37 | 9.79 |
| Val | V | 24 | 6.35 |
| Trp | W | 6 | 1.59 |
| Tyr | Y | 12 | 3.17 |

**Figure 2:**
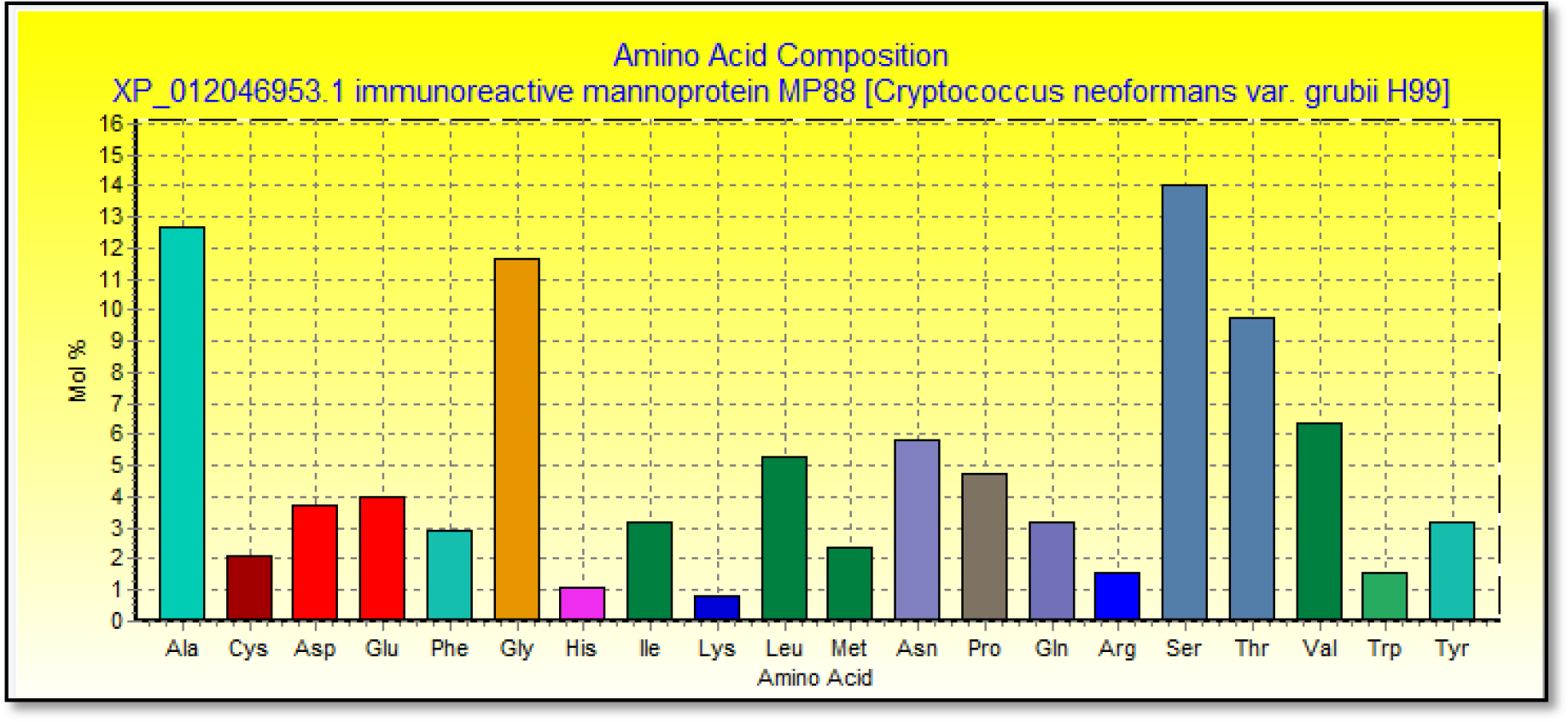
Amino acid composition of mannoprotein MP88, illustrated by BioEdit software version 7.2.5

### Prediction of antigenic B-cell epitopes

To predict linear B cell epitopes which is computationally more feasible, a full length of reference sequence accession number (XP_012046953) was objected to Bepipred Linear Epitope Prediction tool **[36]** at Immune Epitope Database and Analysis Resource (IEDB) with [threshold value=0.350], Bioedit sequence alignment editor used to determined and select conserved epitopes and each conserved epitope was analyze for surface accessibility through Emini Surface Accessibility Prediction tool [Threshold= 1.000] with different window size [37], and Kolaskar & Tongaonkar Antigenicity tool for antigenicity [Threshold= 1.016] with varied window size **[38].**

### T cell epitope prediction

HLA class I and class II T-cell epitope prediction from the conserved sequences by using IEDB tools, where artificial neural network (ANN) method was used to predict 9-mer sequence HLA class I epitopes **[39, 40]** and T cell HLA class II epitopes were identified using *NN-align* method **[41].** All conserved epitopes which bind to alleles at a score less than 500 half-maximal Inhibitory concentrations (IC50) were selected for further analysis.

### Population Coverage

Population Coverage Analysis tool of IEDB was utilized for prediction of population coverage of the whole world. **[42].** This website is designed for calculating epitopes population coverage of different regions based on the distribution of different MHC alleles to which epitopes bind. Predicted MHC I and MHC II epitopes of MP88 were submitted for population coverage tool in IEDB and the calculation option was set to class I and class II.

### Prediction of allergens

An online server was implemented to predict the allergenicity of the selected epitope, named AllerTop 2.0 (http://www.ddg-pharmfac.net/AllerTOP/). Which predict both nonallergens and allergens with a high level of accuracy. The prediction of allergens based on the main physicochemical properties of proteins **[43].**

### Tertiary structure prediction

By using homology modelling technique for protein 3D structure building, RaptorX, (HTTP/www.raptor.uchicago.edu) was used to predict the 3D structure of *C. neoformans var grubii* MP88 by submission of the reference sequence. Obtained 3D protein structure was visualized by UCSF Chimera (version1.8), (http://www.cgl.ucsf.edu/cimera). Missing values in RaptorX PDB file were refined with aid of Modeller software (version 9.19), (https://salilab.org/). Epitopes of B cell, MHC I and MHC II were also visualized within MP88 3D structure each separately.

### Docking

Molecular docking of HLA class 1 peptide was achieved by interacting of 3D structures of MHC I epitope and HLA class 1 molecule using PatchDock server Beta 1.3 version (https://bioinfo3d.cs.tau.ac.il/patchdock). Epitope 3D structures were predicted by De novo peptide structure prediction, PEP-FOLD3 server, while HLA class 1 3D structure was obtained from the PDB database (ID=1b6u). Molecular docking best model was chosen for visualization according to its global energy value.

## Result

### B cell

The reference **Mp88** protein was predicted to have 11 conserved peptides to contain B-cell epitopes. **MP88** protein was analyzed using Bepipred Linear Epitope Prediction that predicts linear epitope; the average binders score of the protein to B cell was 0.571, minimum was −1.956 and 2.816 for a maximum score (Table 2). B cell epitopes illustrate in (figure 5).

**Table 2:**
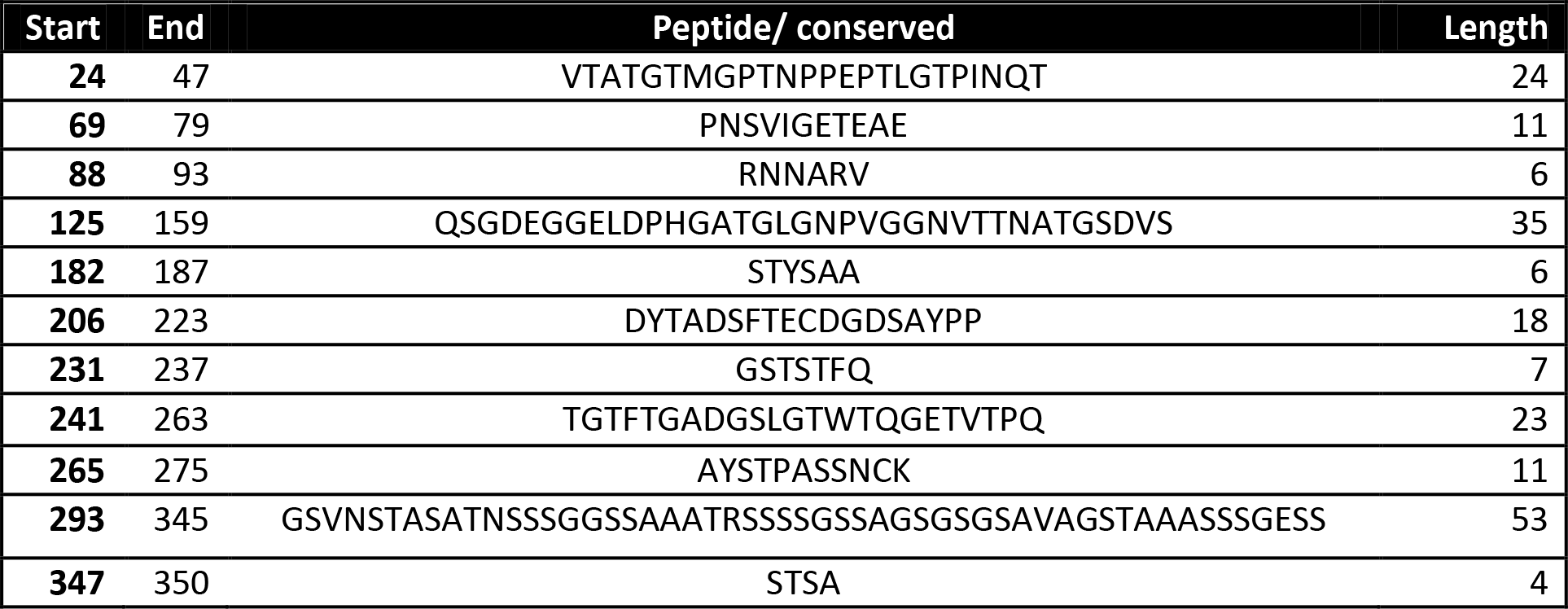
Predicted antigenic B-cell epitopes, 11 antigenic sites were identified from MP88 protein

In Emini surface accessibility prediction; where used for surface accessibility for a potent B cell epitope the average surface accessibility areas of the protein was scored as 1.000, with a maximum of 4.429 and a minimum of 0.065, all values equal or greater than the default threshold 1.000 were potentially in the surface. The result is illustrated in (Table 3) (figure 3).

**Table 3:**
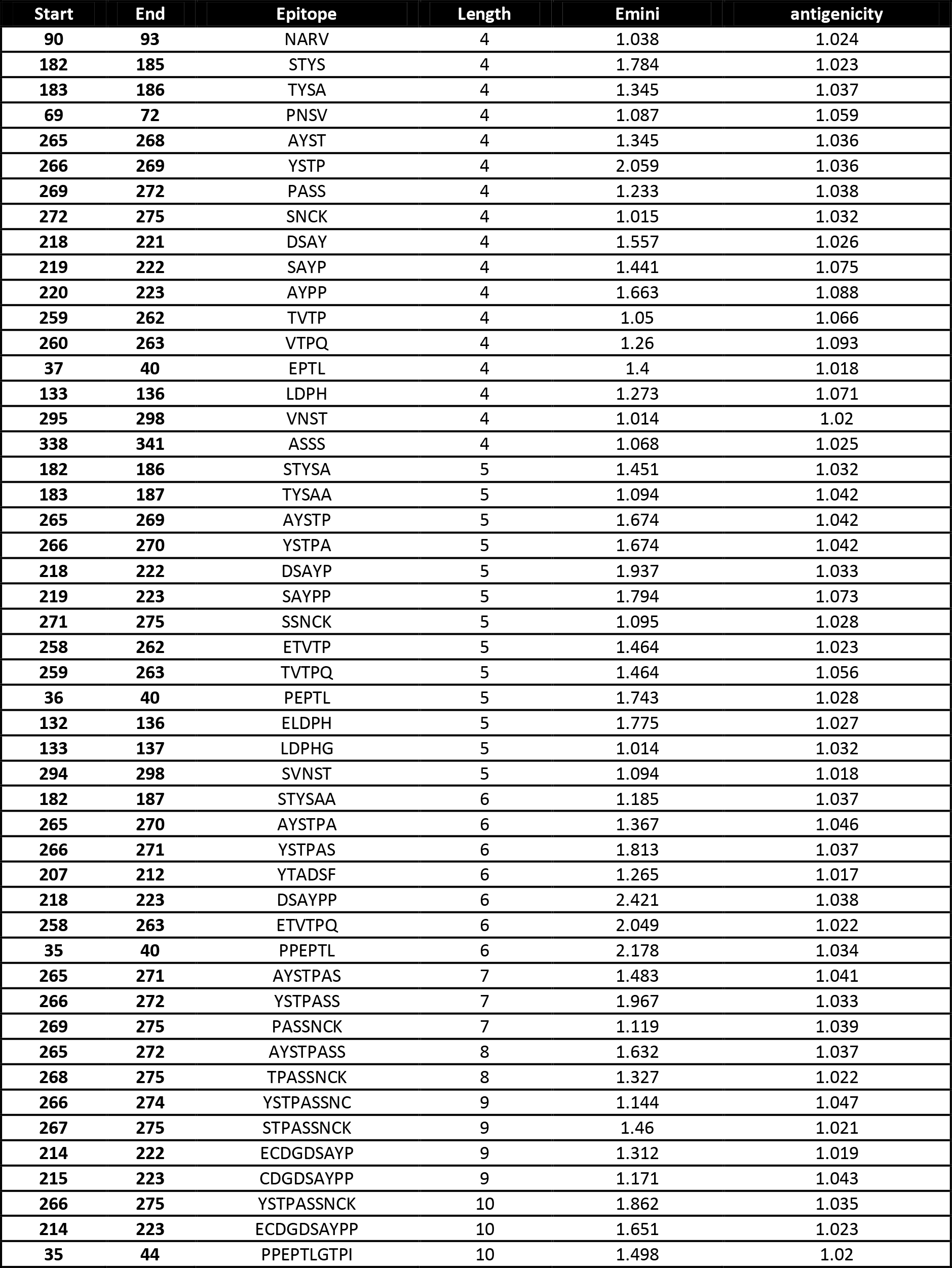
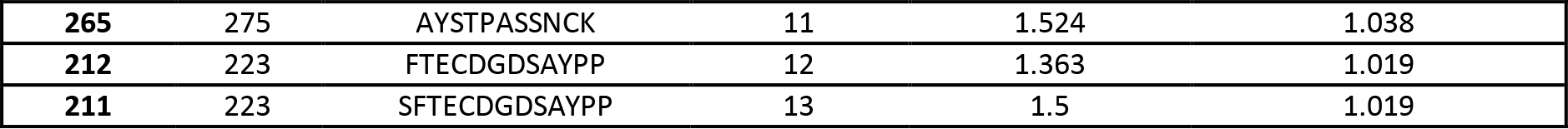
B-cell epitopes prediction:

**Figure 3:**
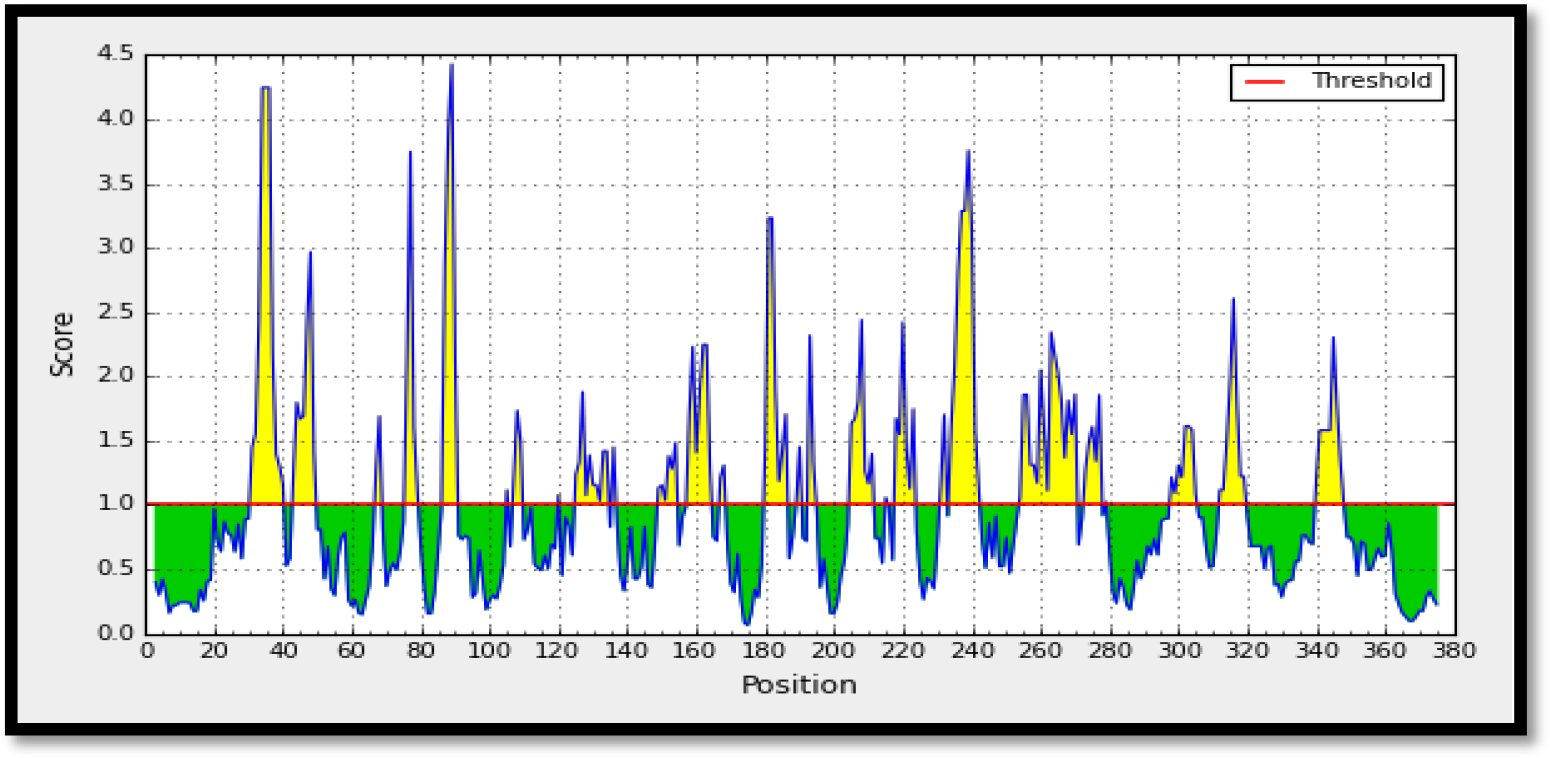
Emini surface accessibility prediction of MP88 protein, the x-axis and y-axis represent the sequence position and surface probability, respectively. The threshold value is 1.000. The regions above the threshold are Surface accessible, shown in yellow.

Kolaskar and Tongaonkar antigenicity prediction method functions on the basis of physiochemical properties of amino acids and abundances in experimentally known epitopes. The average of the antigenicity was 1.016, with a maximum of 1.227 and minimum of 0.883; all values equal to or greater than 1.016 are potential antigenic determinants. The result of all conserved predicted B cell epitopes are shown in (Table 3) and (Figures 4).

**Figure 4:**
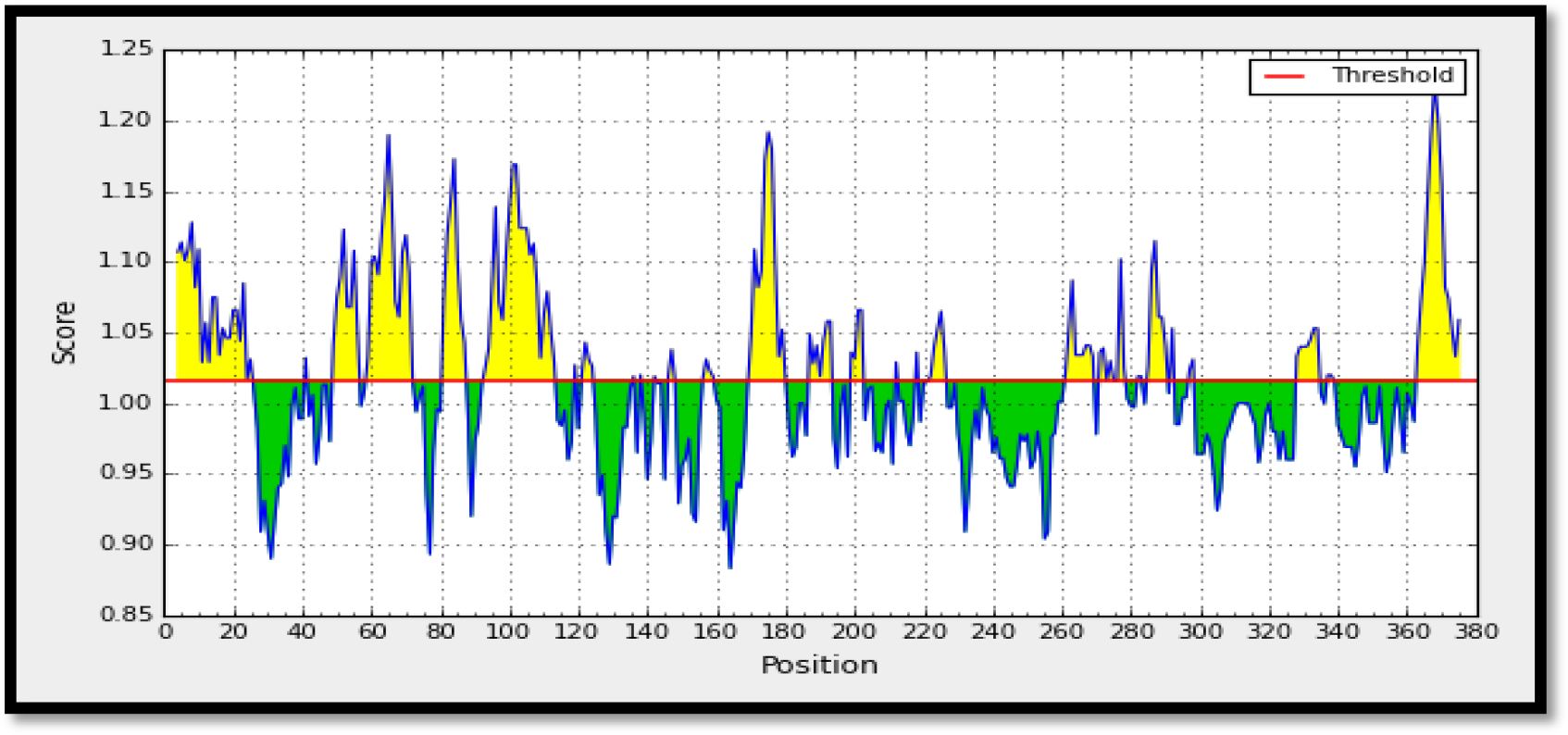
Kolashkar and Tongaonkar antigenicity prediction of MP88 protein, the x-axis and y-axis represent the sequence position and antigenic propensity, respectively. The threshold value is 1.016. The regions above the threshold are antigenic, shown in yellow.

**Figure 5:**
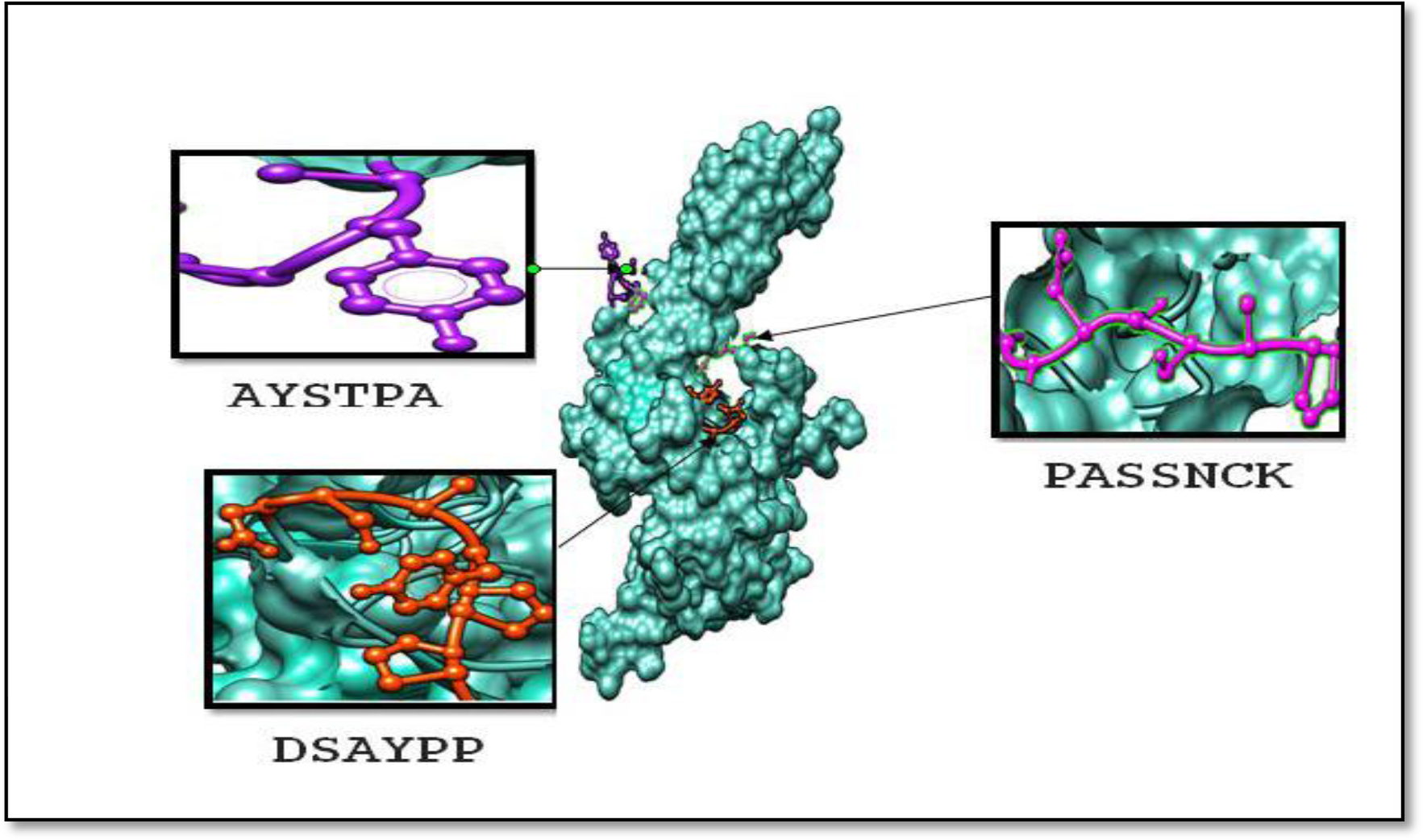
Illustrated of B cell epitopes which visualized with the chimera, version 1.11.2

### T cell

Artificial neural network (ANN) method was utilized to predict the T cell epitopes of the **MP88** reference protein. The most **3** promising conserved epitopes **YMAADQFCL, VSYEEWMNY** and **FQQRYTGTF** were selected according to the lower IC_50_ value. Whereas IC_50_ value less than 500 nM (IC_50_ <500) and this ensured the selection of the MHC-I molecules for which the selected epitopes showed higher affinity. Among the 3 epitopes, the 9-mer epitope **YMAADQFCL** interact with 9 MHC-I alleles and of these 9 alleles HLA-A*02:01 has the best interaction with an epitope ((IC50 = 5.34). These Conserved epitopes can give a more successful immunization (Table 4).

**Table 4:**
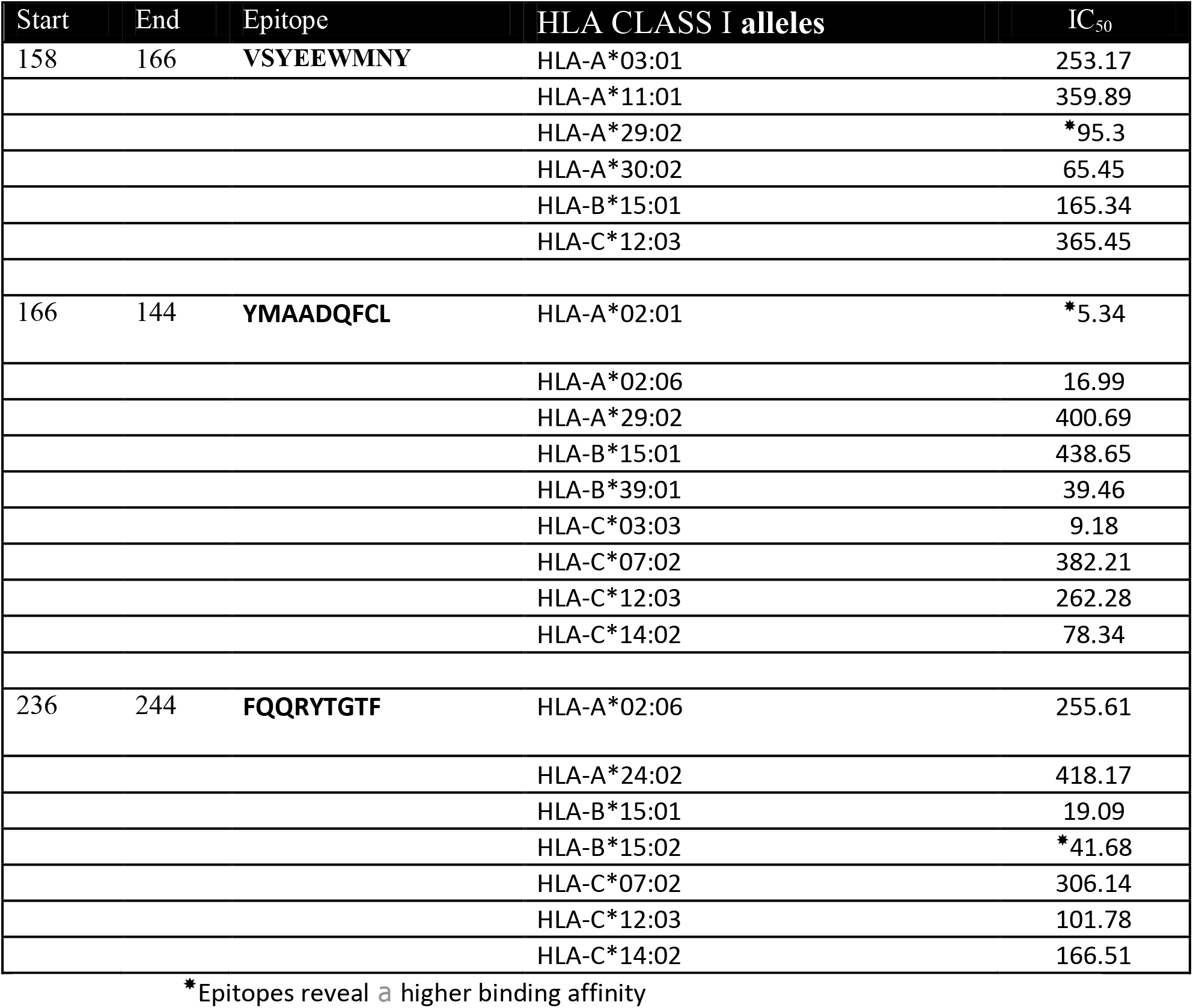
The selected epitopes according to the IC_50_ value predicted by the NNA method

The 3 epitopes were objected into AllerTop 2.0 server to predict allergenicity, **VSYEEWMNY** and **FQQRYTGTF** they are non-allergen while **YMAADQFCL** was an allergen.

For MHC class II peptide binding prediction, we used NN-align prediction tool, where **YARLLSLNA, ISYGTAMAV** and **INQTSYARL** represent the most Three highly binding affinity core epitopes with IC_50_ value <1000. Among the 3 core epitope **INQTSYARL** was found to interact with 14 MHC-II alleles while **ISYGTAMAV** and **YARLLSLNA** similarly interact with 12 MHC-II alleles (Table 5). For allergenicity prediction the **ISYGTAMAV, INQTSYARL** they are non-allergen and **YARLLSLNA** was an allergen.

**Table 5:**
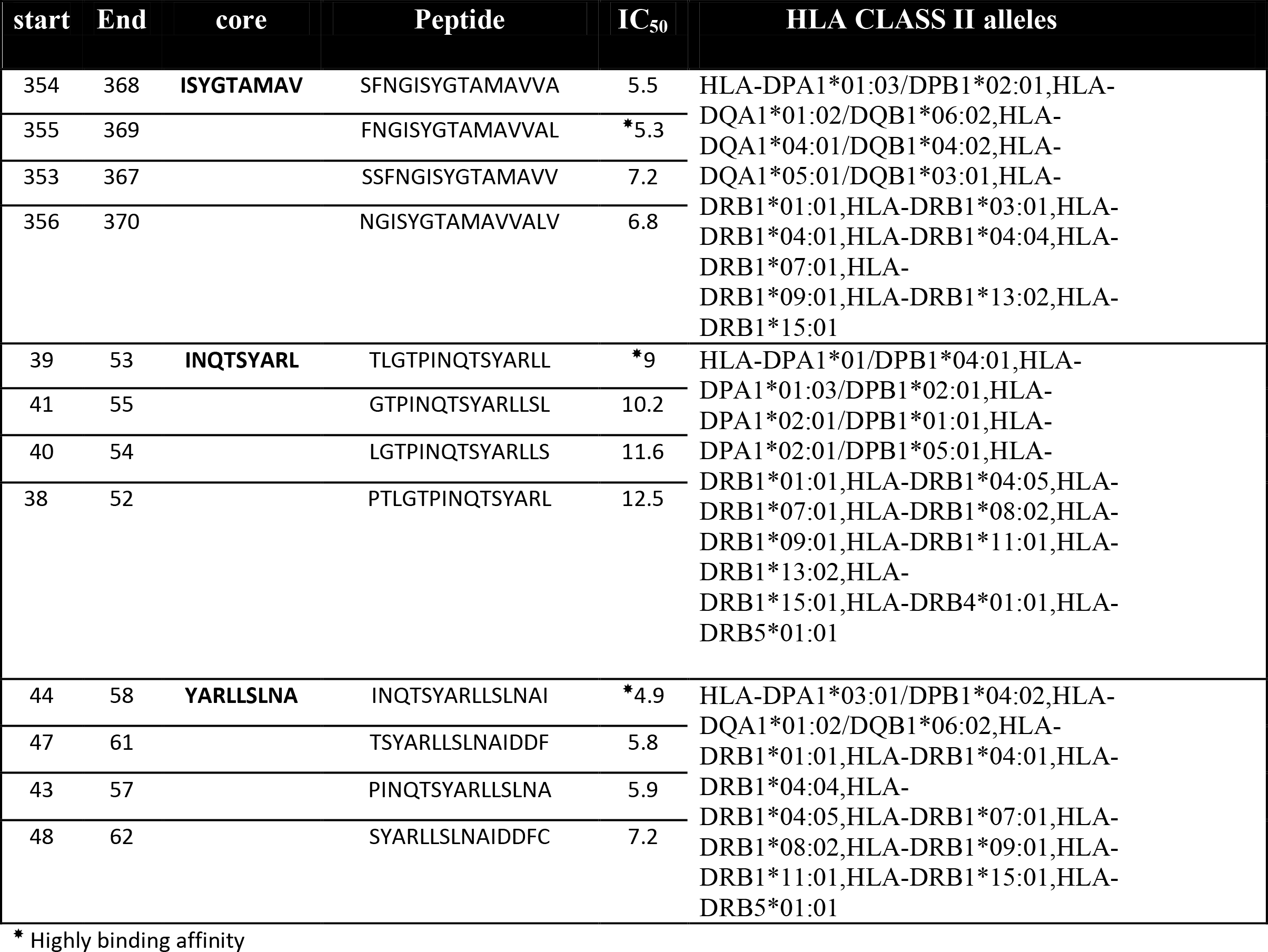
Candidates of the T-cell epitope for MHC class II.

### Analysis of the Population Coverage

Population coverage was taken to cover maximum possible populations. In MHC class I, **YMAADQFCL, VSYEEWMNY** and **FQQRYTGTF** gave a high percentage against the whole world population by IEDB population coverage tool. The population coverage of **YMAADQFCL** exhibits a higher percentage among the world (69.75%) while **VSYEEWMNY** and **FQQRYTGTF** display (58.78%, 54.11%) respectively. The average population coverage was **93.01%** (**figure 6**).

**Figure 6:**
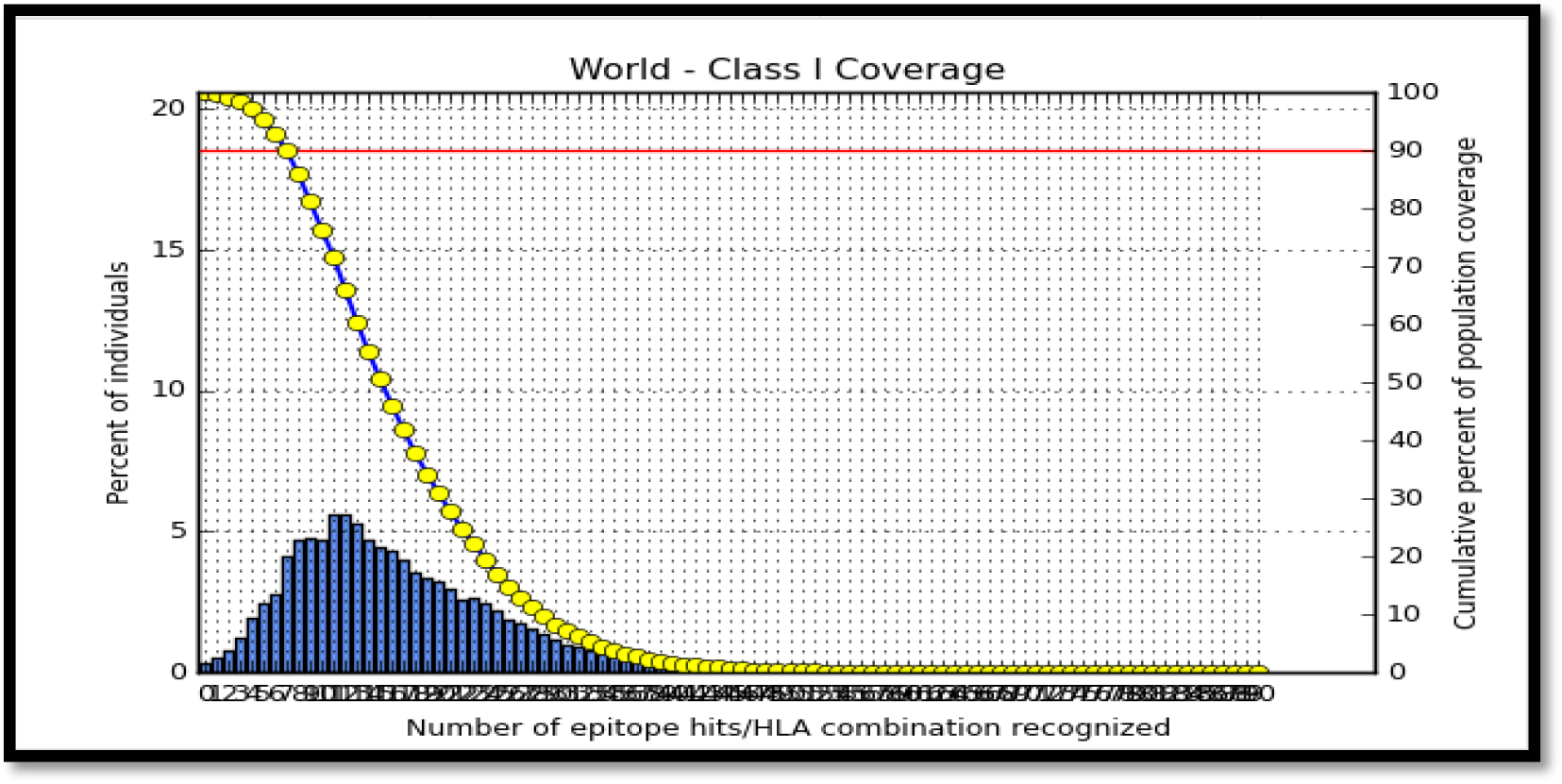
illustrated of Population coverage of MHC class I

In MHC-II, **ISYGTAMAV** epitope reveal higher percentage (74.39%) while **YARLLSLNA** and **INQTSYARL** showed (69.46%, 63.94%) respectively. The everage population coverage was (81.94%).on the other hand following alleles (HLA-DQA1*05:01/DQB1*03:01, HLA-DQA1*01:02/DQB1*06:02, HLA-DPA1*01:03/DPB1*02:01, HLA-DPA1*01/DPB1*04:01, HLA-DPA1*02:01/DPB1*05:01, HLA-DRB4*01:01, HLA-DRB5*01:01, HLA-DQA1*04:01/DQB1*04:02, HLA-DPA1*03:01/DPB1*04:02, HLA-DPA1*02:01/DPB1*01:01) are not available on the IEDB and therefore not included in the calculation(Figure 7)

**Figure 7:**
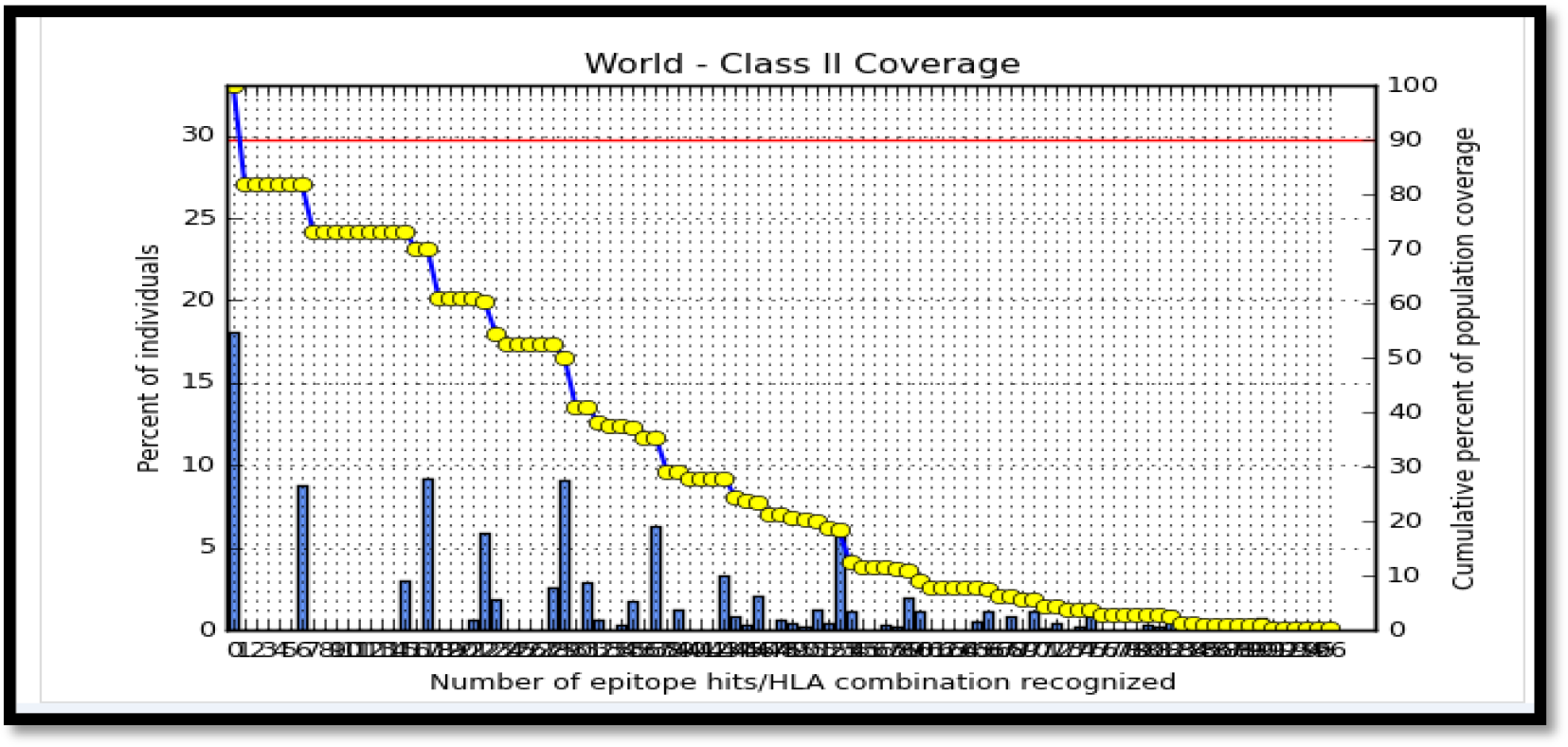
illustrated of Population coverage of MHC-II

### Molecular docking analysis

**HLA-A*02:01** has the best interaction with the **YMAADQFCL,** therefore was chosen as a model for molecular docking of HLA class 1 peptide. **HLA-A*02:01** has 3D structure available in PDB database (ID= 1b6u). The 3D structures of epitopes were predicted using PEP-FOLD [**44, 45].** (**Figure 8**).

**Figure 8:**
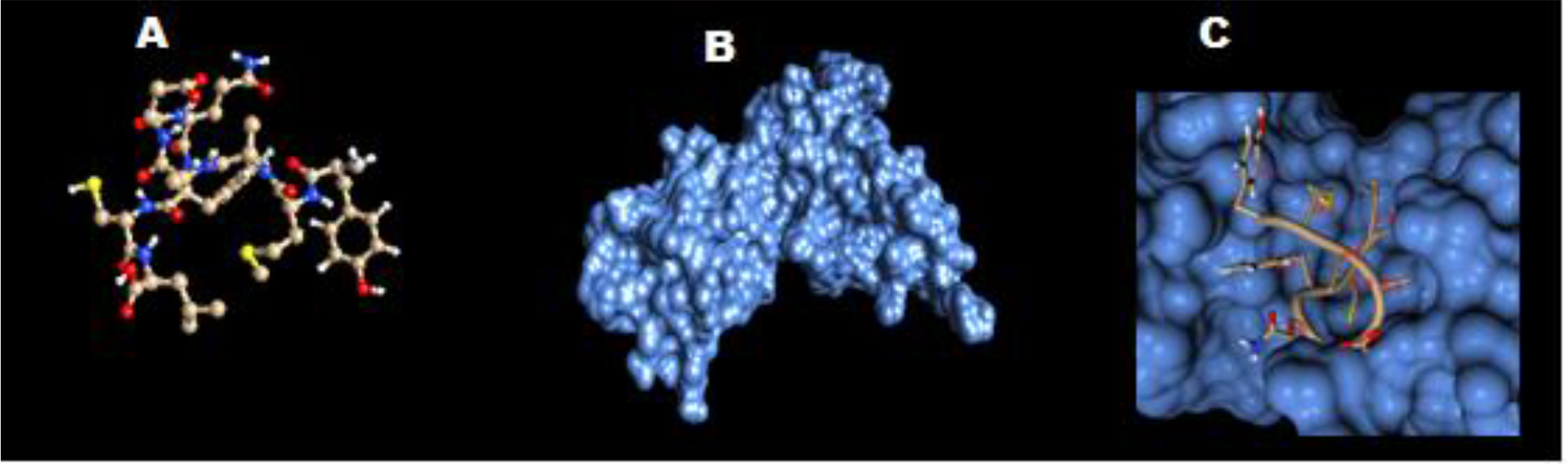
(A) Structure of predicted epitope “YMAADQFCL”, (B) Structure of predicted HLA-A*02:01” (C) Docking of YMAADQFCL with HLA-A*02:01, docking interaction was visualized with the chimera, version 1.11.2

## Discussion

In the present study, we successfully designed a peptide vaccine for ***C. neoformans*** against its immunogenic mannoprotien (**MP88**) by immunoinformatics tools. Predicted peptide vaccine characterized by easy production, stimulating effective immune response and no potential infection possibilities [35].

B-cell epitopes are regions in a pathogenic antigen recognized by B cell receptors or antibody to induce specific humeral response [46]. Linear B cell epitopes of MP88 were achieved by Bepipred linear epitope prediction tool in (IEDB). Fully conserved epitopes were testified for surface accessibility and antigenicity by aid of Emini surface accessibility prediction method and Kolaskar and Tongaonkar antigenicity method respectively. According to their high scores in Emini surface accessibility and antigenicity tests epitopes 265 **AYSTPA** 270, 265 **AYSTPAS**271, 269**PASSNCK** 275 and 218**DSAYPP** 223 were approved as promising B cell epitopes.

T cell CD8+ recognize antigen of a pathogen only when combined with MHC I molecules and trigger a cytotoxic response [47]. ANN method precisely determined three promising T cell epitopes within MP88; 166 **YMAADQFCL**174, 158**VSYEEWMNY** 166and 236 **FQQRYTGTF** 244. These epitopes could potentially induce CD8+ T cell immune response when interacting strongly with MCH I alleles such as YMAADQFCL/HLA-A*02:01, FQQRYTGTF/HLA-B*15:01 and VSYEEWMNY/HLA-A*29:02. These promising epitopes interact with the most predominant HLA class II allele in human like HLA-A*02:01 [49, 50], in same time will provide protection against Cryptococcal infection for 93% of world population in case if used in forward vaccine.

T cell CD4+ or chestrates the adaptive immune response in human body [48]. Specific interact with high binding affinity epitope / HLA allele class II molecule unleash protective and specific adaptive immune response. The obtained T cell CD4+ epitopes **YARLLSLNA, ISYGTAMAV** and **INQTSYARL** from NN-align prediction tool represent the most effective epitopes which can induce specific and potent adaptive immune response against Cryptococcal MP88. Immunization with these three epitopes will cover about 81% of world population.

Cryptococcosis is the most fungal infection in immunocompromised individuals especially who’s with HIV infection, cancer or solid organ transplantation patients. Very serious complications starting from severe pulmonary infection until fatal meningoencephalitis were reported [51–54]. High cost of Cryptococcosis treatment and some clinical consequences like toxicity and antifungal resistance [14, 55] lead to exhibiting a strong prolonged preventive approach. Until now there is no applied accredited vaccination for Cryptococcosis.

presumably, Killed vaccine against ***C. neoformans*** classified as unsuccessful ones, while live attenuated vaccine have been shown inducing protective immunity in immunocompetent animals but not practical for immunocompromised hosts[51].

Previously, GXM known as poor immunogenic antigen as solo component of cryptococcal capsule. Multiple studies shown that immunization by GXM based vaccine even conjugated with a protein or Tetanus Toxoid (GXM-TT), fail to induce specific protective immune response and often act as deleterious factor [51, 56].

Now, these proposed vaccines still under experimental trials. A Study done by Upadhya and his colleagues revealed that heat-killed chitosan of Cryptococcal cell wall vaccine could develop robust protective immunity against virulent strains of ***C. neoformans*** in mice which represent as potential vaccine candidate [55]. *Specht etal* in their study reported experimental vaccine based on alkaline extract from mutant Cryptococcal strains packaged in to glucan particles (GP) could provide significant protection in mice, but identifying the potent molecule is still in search [57]. In other hand GXM mimotope based vaccine is likely consider as promising protective epitopes [56, 57], but the difficulty lie in focusing on response of single epitope could enhance the risk of escape variants [56].

These vaccinexperiments are likely costly, beside they don’t take along the genetic diversity of ***C. neoformans*** antigensand the world population coverage of the expected vaccine which it was considered in our study.

Our study met many reports elicit mannoprotiens the minor component of ***C. neoformans*** capsule as a strong potent vaccine candidate [51, 56, 58], which could provide a protective T h 1 type immune response versus ***C.neoformans***infection. Vaccination of mice models with MPs/Ribi adjuvant demonstrate an apparent T cell immune response but not antibody response [59].

This is the first in silico approach to predict***C. neoformans*** epitope vaccine based on immuoninformatics tools, which will facilitate manufacturing a specific, protective vaccine for immunocompromised patients around world combined with reducing in practical efforts. Successful peptide vaccine should compose of B cell and T cell epitopes in one vaccine. Whereas peptide vaccine is poorly immunogenic when used alone [60], in coming studies were needed to select suitable delivery system or adjuvant and strategies which could improve*in vivo* T cell and antibody immune responses.

Despite of the significant results of our findings, little objections were noticed. There is a need to expand number of Cryptococcal MP88 protein (378 aa) retrieved sequences from all parts of the world when it is available in global protein data base to reduce margin of error. Calculations of some HLA class II alleles/epitopes for population coverage were missed, that influence in generalizing of results.

